# A G1 sizer mechanism coordinates growth and division in the mouse epidermis

**DOI:** 10.1101/754424

**Authors:** Shicong Xie, Jan M. Skotheim

**Affiliations:** Department of Biology, Stanford University, Stanford, CA 94305

## Abstract

Cell size homeostasis is often achieved by coupling cell cycle progression to cell growth. Studies of cell size homeostasis in single-celled bacteria and yeast have observed several distinct phenomena. Growth can be coupled to division through a range of mechanisms, including a ‘sizer’, wherein cells of varying birth size divide at similar final sizes [1–3], and an ‘adder’, wherein cells increase in size a fixed amount per cell cycle [4–6]. Importantly, intermediate control mechanisms are observed, and even the same organism can exhibit distinct control phenomena depending on growth conditions [2,7,8]. While studying unicellular organisms in laboratory conditions may give insight into their growth control in the wild, this is less apparent for studies of mammalian cells growing outside the organism. Sizer, adder, and intermediate mechanisms have been observed *in vitro* [9–12], but it is unclear how these diverse size homeostasis phenomena relate to mammalian cell proliferation *in vivo*. To address this gap, we analyzed time-lapse images of the mouse epidermis taken over one week during normal tissue turnover [13]. We quantified the 3D volume growth and cell cycle progression of single cells within the mouse skin. In dividing epidermal stem cells, we found that cell growth is coupled to division through a sizer mechanism operating largely in the G1 phase. Thus, while the majority of tissue culture studies to-date identified adder mechanisms, our analysis demonstrates that sizer mechanisms are important *in vivo* and highlights the need to determine their underlying molecular origin.

## Results

### Measuring cell volume growth in epidermal stem cells during normal tissue turnover

To determine which cell size homeostasis mechanism operates *in vivo*, we examined epidermal stem cells growing and dividing during normal tissue turnover. Mouse skin is an ideal system to study *in vivo* cell size control because it has a high frequency of cell divisions in healthy adults [14]. The epidermis is a multilayered epithelium with suprabasal layers of differentiated keratinocytes residing above a basal layer of stem cells (Figure 1A). The basal layer epidermal stem cells are the only source of new cells during normal tissue turnover [15]. As they proliferate, epidermal stem cells can either self-renew and remain in the basal layer or differentiate into the suprabasal layers, where they eventually shed.

**Figure 1.**
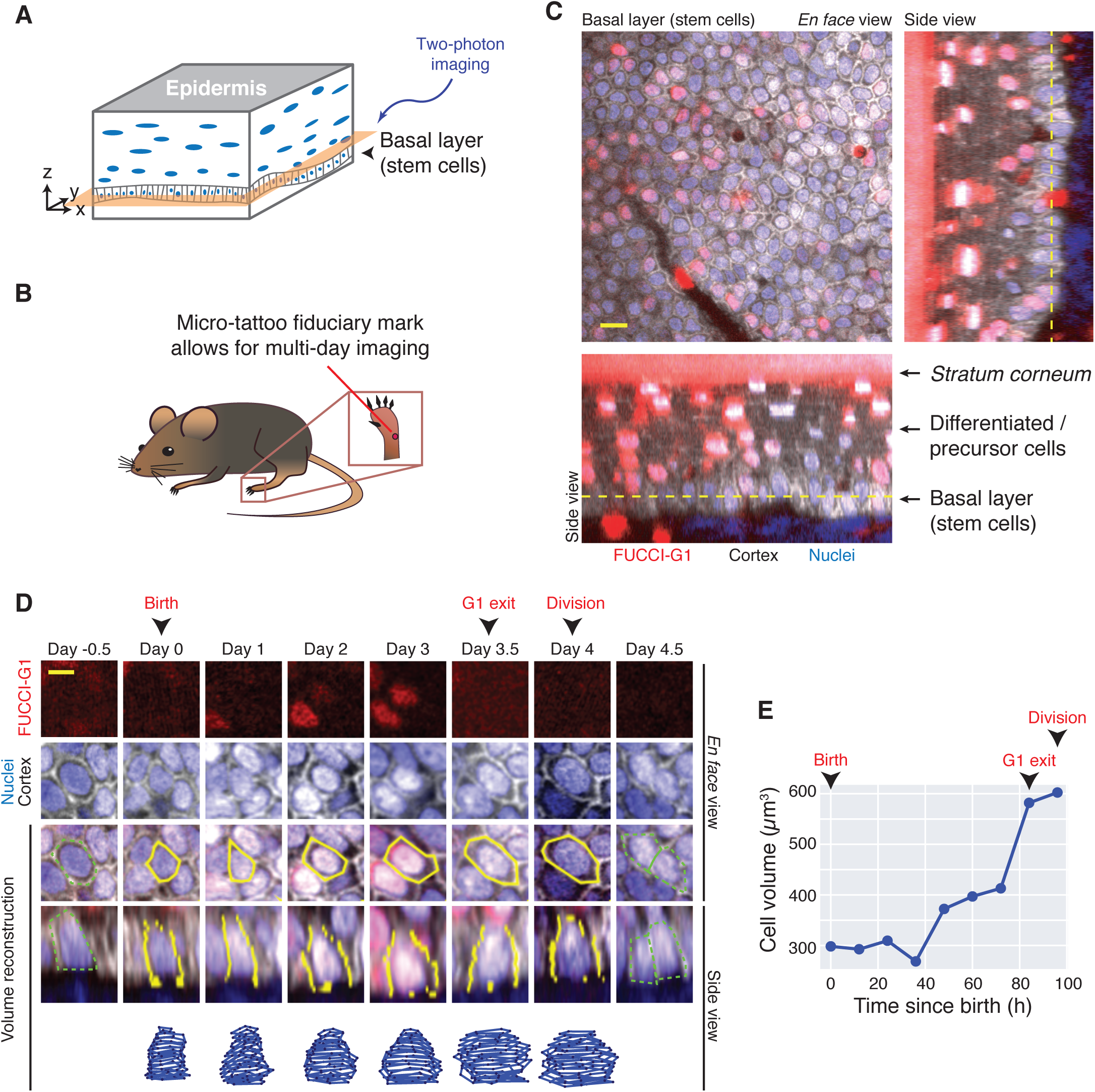
Quantifying cell volume and cell cycle phase of epidermal stem cells growing *in vivo*. A. Schematic of mouse non-hairy plantar skin. The epidermal tissue is a stratified epithelium, where stem cells reside in the basal layer. During tissue homeostasis, the basal layer is the only population of cycling cells. B. Dataset from Mesa, Kawaguchi, & Cockburn, *et al*. (2018) [13]. Movies of cells in the mouse hind paw skin were generated by re-visiting a micro-tattooed region with two-photon microscopy every 12h for up to 7 days. C. Epidermal stem cells in a mouse expressing reporters for G1 phase (red, *CMV-mKO2-hCdt1(30/120)*), cell cortex (gray, *K14-actin-GFP*), and nucleus (blue, *K14-H2B-Cerulean*). The *en face* view as well as two side views are shown. Dotted lines denote the z-position of the *en face* view shown. Scale bar is 10 µm. D. Example of the volume reconstruction of a single epidermal stem cell. The FUCCI-G1 reporter is shown in red, the nucleus in blue, and the actin cortex in gray. The manually segmented cell outlines are shown in yellow on top of the merged images showing *en face* and side views. The reconstructed 3D shape is shown in the bottom track. The cell cycle landmarks birth, G1 exit, and division are annotated. Note the presence of the parent cell at day −0.5 and daughter cells at day 4.5 outlined in dotted green. Scale bar is 5 µm. See also Supplemental Figure S1. E. The volume growth curve for the cell shown in (D). Birth, G1 exit, and division are marked.

To measure cell growth and division *in vivo*, we analyzed images previously acquired by Mesa, Kawaguchi, and Cockburn, *et al.* (2018) [13]. Skin regions on the mouse hindpaw were micro-tattooed, which served as fiducial marks allowing re-visiting of the same regions over time (Figure 1B). Cells in the same region were imaged with two-photon microscopy every 12 hours for up to 7 days (Figure 1C). Since mice were not wounded during the experiment and were allowed to return to normal activity between imaging, these movies capture stem cell growth dynamics during unperturbed tissue turnover.

To quantify cell volume, we reconstructed 3D cell shapes by manually segmenting movies from mice expressing Histone-2B-Cerulean (nuclear reporter), Actin-GFP (cortex reporter), and FUCCI-G1 mKO2-hCdt1(30/120) (G1 reporter) (Figure 1D-E, Supplemental Figure S1). We could track cells over an entire cell cycle and use the FUCCI-G1 reporter to distinguish G1 from S/G2/M phases. We restricted our analysis to dividing stem cells within the basal layer. Similar 4D reconstruction strategies have been used to quantify *in vivo* epithelial cell volume and shape change in *Drosophila* and *Arabidopsis* [16, 17].

We measured cell volume growth over entire cell cycles for 197 cells from 3 independent tissue regions in 2 different mice (Figure 2A, Supplemental Figure S2A-C; **Movie S1-2**). Cells on average cycled every 71 ± 21h (std), with the majority of time spent in G1 phase (Figure 2B-D). This estimate is consistent with independent estimates of hindpaw epidermis cell cycle durations [18]. Notably, cell cycles *in vivo* are much longer than cell cycles *in vitro*, where cell lines typically divide once a day. To assess the error in our 3D segmentation, we fitted our data to smoothing splines, since cell volume is expected to increase gradually over time (Supplemental Figure S3A-F). Over 90% of fitting residuals for our spline fits are smaller than 50µm^3^ in magnitude, and the average residual is <10% of the estimated volume signal (Supplemental Figure S3G-H). We also tested the intra-user variability in 3D segmentation by performing independent repeat measurements and found that the average intra-user error is similar in magnitude (9.1± 8.6%, std) (Supplemental Figure S3I-J). Next, we compared the volume growth curves with cross-sectional area growth curves, a common cell size proxy in epithelial cells (Supplemental Figure S4A-B). Area growth curves, while correlated with volume growth (Supplemental Figure S4C), are significantly noisier with larger relative residuals when fit to smoothing splines (Supplemental Figure S4D). These analyses suggest that our volume growth data have low error and better reflect cell size than cross-sectional cell area measurements. To reduce error in subsequent analyses, we present spline-smoothed data in the main text. Similar analyses on raw data yielded qualitatively similar conclusions and are shown in the supplemental figures.

**Figure 2.**
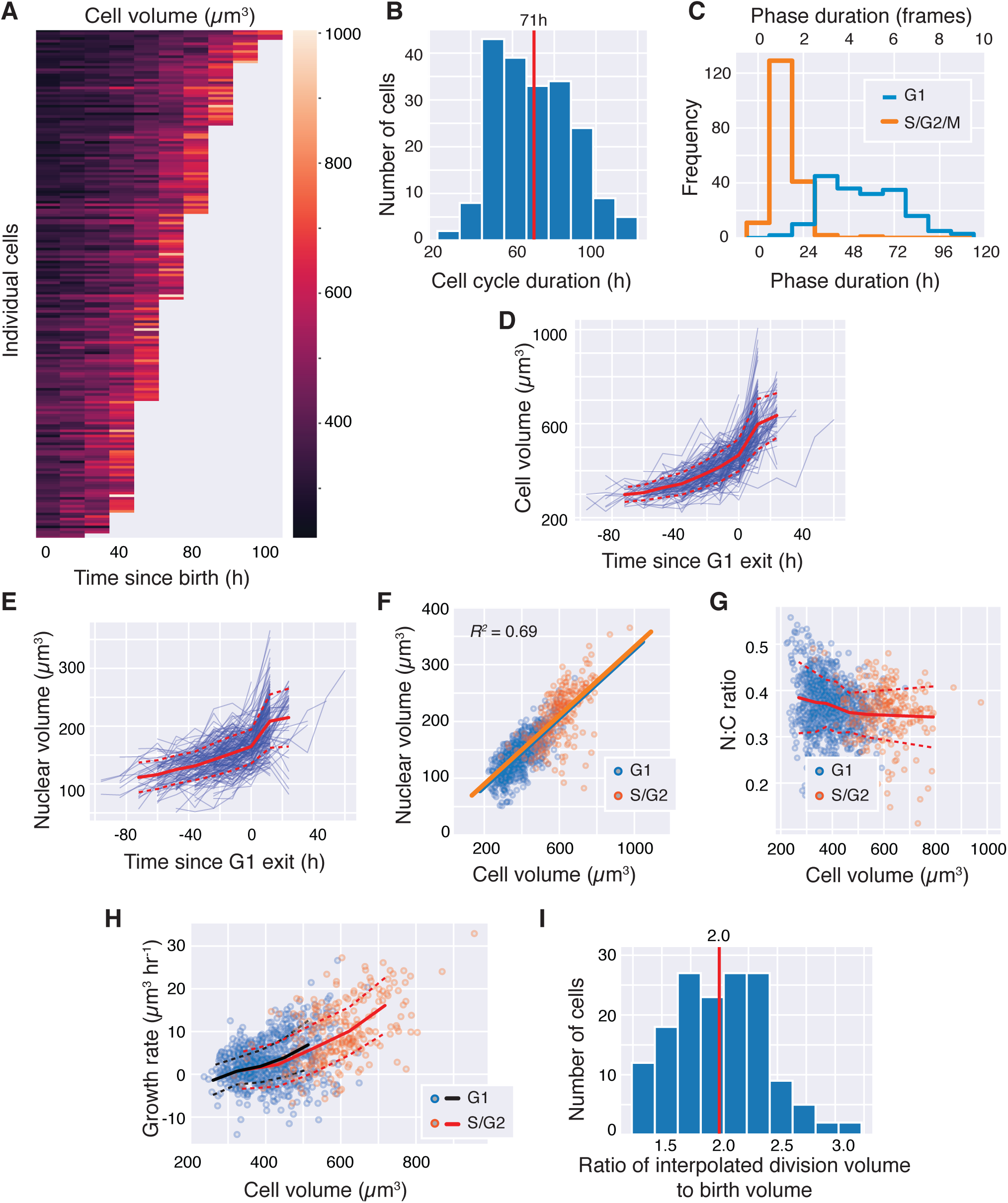
Epidermal stem cells grow faster than linearly and maintain a constant nuclear to cytoplasmic ratio. A. Heatmap of volume growth curves from epidermal stem cells (*N* = 197; 3 independent regions from 2 mice). Growth curves are sorted by increasing length. B. The distribution of cell cycle durations. C. The distribution of the duration of G1 (blue) and S/G2/M (orange) phases. D. Volume growth curves aligned by the time of G1 exit. E. Nuclear volume growth curves aligned by the time of G1 exit. F. The correlation between cell and nuclear volumes (*N* = 1,159). Straight lines show linear regression for G1 cells (blue) or S/G2 cells (orange). G. The nuclear-to-cytoplasmic volume ratio (N:C ratio) for cells of varying volumes. Blue dots denote G1 cells, orange dots denote S/G2 cells. H. The cell volume growth rate plotted as a function of cell volume (*N* = 946). The binned mean values are shown for G1 (black) and S/G2 (orange) cells. Dotted lines denote standard deviations corresponding to their color. I. The distribution of the ratio of the interpolated division volume to the birth volume (*N* = 167). Means or binned mean values are shown in red solid lines. Dotted lines denote standard deviations. See also Supplemental Figure S2 for data broken down by imaging region and mouse.

### Nuclear-to-cytoplasm ratio remains constant throughout the cell cycle

Nuclear volume has been shown to be proportional to cell volume (constant N:C ratio) in different cell lines. While the N:C ratio may vary across cells of different types and/or species, within the same cell type it is typically independent of cell size and cell cycle phase [19]. To test if this remains true *in vivo*, we quantified nuclear volume by segmenting the nucleus using labeled histone H2B-Cerulean (Supplemental Figure S5). We observed that nuclear volume grows during cell growth, and that the N:C ratio remains relatively constant throughout both G1 and S/G2 phases (Figure 2E-G; Supplemental Figure S6A-B). In bivariate regression analysis using cell volume or cell phase to predict nuclear volume, cell volume is a clearly significant predictor (*P* < 10^-10^) while cell phase is not (*P* > 0.05). Thus, the N:C ratio remains relatively constant throughout the cell cycle *in vivo*.

### Cells grow faster than linearly *in vivo*

Next, we examined the volume growth dynamics of epidermal stem cells *in vivo*. Currently, there is conflicting evidence for how mammalian cells grow. *In vitro*, cells have been reported to grow linearly (accumulating a constant amount of volume per unit time), exponentially (accumulating volume in proportion to their current volume), or with intermediate dynamics [10,12,20–22]. However, there is no comparable data in mammalian tissues *in vivo*, where single-cell growth rates are especially difficult to measure. We quantified the absolute growth rate as a function of cell volume by taking the difference in cell volume between one time-point and its previous time-point, normalized by the framerate. Stem cells grow faster the larger they become, leading to supra-linear growth (Figure 2H). Supporting this, volume growth curves better fit exponential compared to linear models (Supplemental Figure S3G). This supra-linear relation is also present in the raw data (Supplemental Figure S6C). Furthermore, there is little difference in growth rates between G1 and S/G2/M cells of the same size. In bivariate regression analysis where cell volume and cell cycle phase were used to predict growth rate, cell volume is a significant predictor (*P* < 10^-10^), whereas cell cycle phase is not (*P* > 0.9). Similarly, if we break G1 into early G1 (first 3 frames) and the rest of G1, cell volume significantly predicts growth rate in G1 (*P* < 10^-10^) while early or late G1 phase does not (*P* > 0.9). Taken together, these results suggest that size rather than cell cycle phase is the dominant factor determining cell growth rate. In both G1 and S/G2/M phases, growth dynamics is faster than linear.

### Estimating the final cell division size

Since cells grow faster when they are larger, we hypothesized that the 12h sampling rate may lead to a significant underestimation of the final division size of the cell if we take the division size at the final frame before daughter cells appear. We therefore estimated division size by averaging the size at the last frame with the sum of the sizes of the daughter cells in the subsequent frame (Supplemental Figure S7A-C). Stem cell divisions result in daughters of high symmetry (Supplemental Figure S7D). We observed that the interpolated division size is on average 35 ± 57 µm^3^ (std) larger than the sampled final size before division (Supplemental Figure S7E-F). Importantly, the interpolated volume is likely a better estimate of the division volume because it is closer to twice the average birth volume (1.9-fold vs 2.0-fold; *P* < 10^-13^, one-sided paired T-test) (Figure 2I; Supplemental Figure S7G).

### Evidence for swelling during mitosis

Recent studies showed that cells *in vitro* rapidly swell up to 20% in volume when entering mitosis, and this swelling is quickly lost during anaphase [23, 24]. This osmotically-driven swelling allows cells to push away their neighbors and achieve a spherical geometry during mitosis, which is thought to support spindle formation and accurate chromosome segregation [25]. Interestingly, we have 11 growth curves where the final frame before division captured cells in mitosis as evident in their spherical shape and loss of nuclear envelope (Supplemental Figure S7H). Consistent with the mitotic swelling model, we found that the volume of mitotic cells is larger than the sum of the two daughter volumes following division (Supplemental Figure S7I).

### G1 and S/G2/M durations adjust to birth size variation

To determine how cell size is controlled in epidermal stem cells, we first examined the relationship between cell size and cell cycle phase durations. Cells born smaller are much more likely to spend longer in G1, suggesting that the G1/S transition is important for cell size control (Figure 3A; *R* = −0.54, *P* < 10^-15^). Cells born in the smallest size bin (233-291 µm^3^) on average spend 70h ± 19 (std) in G1, whereas cells born in the largest bin (398-496 µm^3^) spend 40h ± 13 in G1. We note that cells exiting G1 with smaller volumes also spend on average longer in S/G2/M (Figure 3B; *R* = −0.47, *P* < 10^-10^). Cells exiting G1 in the smallest size bin (290-432 µm^3^) spend 20h ± 12 in S/G2/M, while cells exiting G1 in the largest size bin (530-671 µm^3^) spend 11h ± 4 in S/G2/M. However, there is poor sampling of the S/G2/M phases which only last 1-2 movie frames (Figure 2C). This low sampling may introduce artifacts in the S/G2/M correlations (see **Discussion**; Supplemental Figure S8). Notably, the coefficient of variation (CV) of the G1 exit volume is smaller than the CV of cell size at birth or at cell division (Supplementary Figure S9; *P* < 0.05, bootstrap test). Taken together, these data suggest that there may be size control occurring at the G1/S transition.

**Figure 3.**
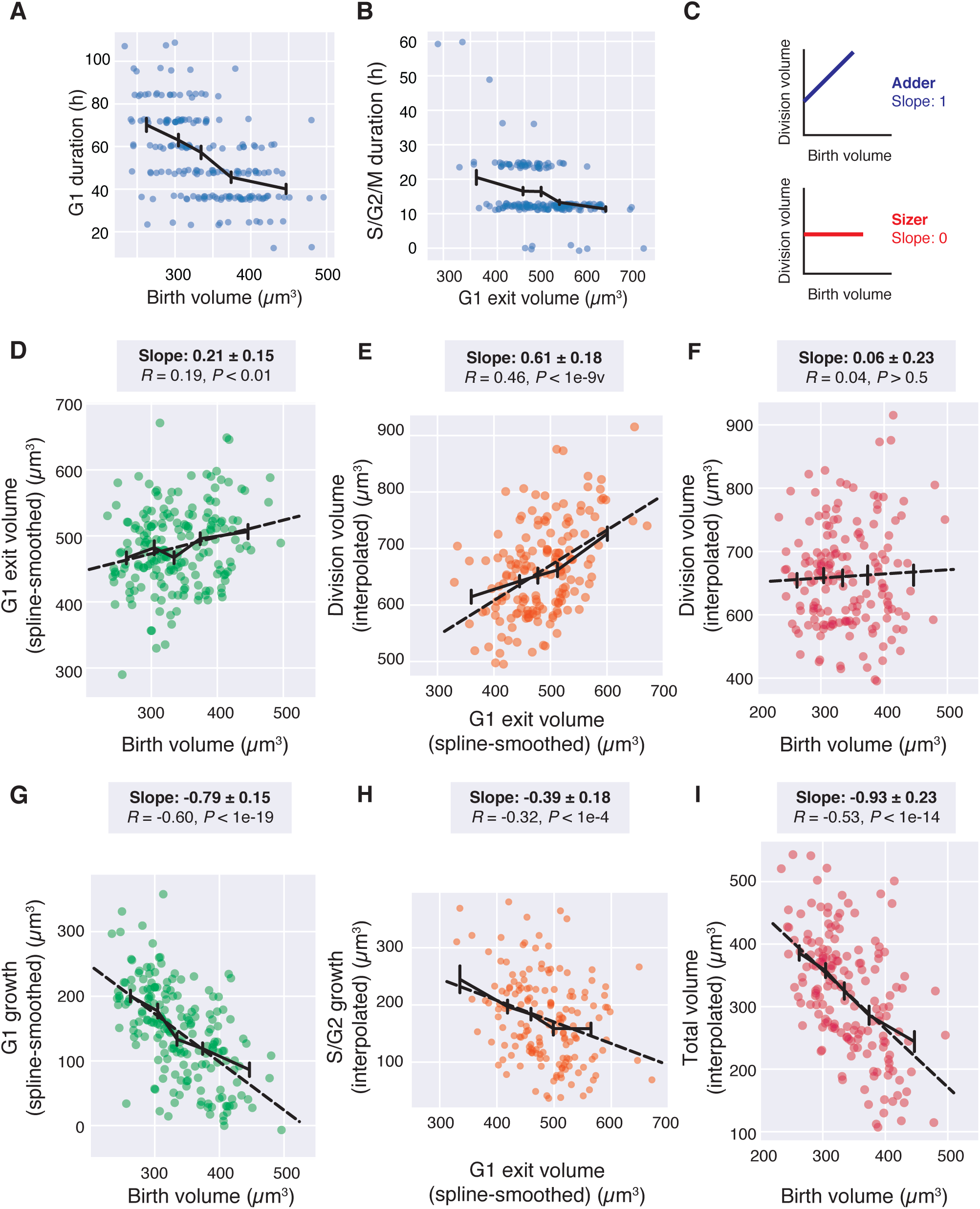
Epidermal stem cells exhibit sizers. A. Cell birth volume is plotted against the duration of G1 phase (*N* = 197). Small jitter was added in the plot to avoid overlapping data so all points can be seen. B. The volume at which cells exit G1 is plotted against with the total duration of S/G2/M phases (*N* = 197). C. Schematic of cell size control correlations. For the adder model, the linear regression slope between birth and division volume is 1. For the sizer model, the slope is 0. D. The cell birth volume is plotted against the G1 exit volume (*N* = 197). E. The G1 exit volume is plotted against the volume at division (*N* = 167). F. The birth volume is plotted against the volume at division (*N* = 167). G-I. The amount of volume grown during the respective phases are plotted against the cell volume at the beginning of the indicated phase (G, H) or the entire cell cycle (I). Solid black lines show the binned mean ± SEM. For D-I, dotted black lines show the linear regression. Linear regression slopes are reported above the plots with 95% confidence intervals. *R* values are Pearson’s correlations, with *P*-values reported against the null-hypothesis of *R* = 0. See also Supplemental Figure S10.

### Epidermal stem cells grow as sizers

That smaller cells spend more time in G1 phase suggested that they would be able to compensate for their small initial size by growing more during their cell cycle. If the G1/S transition were a ‘sizer’, the linear regression slope (*m*) between birth size and G1 exit size would be *m =* 0. Conversely, if G1/S were an adder, then the slope would be *m =* 1 (Figure 3C). We find that for epidermal stem cells, the slope of the linear regression between birth volume and G1 exit volume is *m* = 0.21 ± 0.15 (95% confidence interval), indicating a near-sizer mechanism coupling cell birth size to G1 exit (Figure 3D). During S/G2/M, the correlation between G1 exit volume and growth during S/G2/M is intermediate between a sizer and adder with *m* = 0.61 ± 0.18 (Figure 3E). Over the entire cell cycle, the slope between birth volume and division volume is *m* = 0.06 ± 0.23, which corresponds to an almost perfect sizer (Figure 3F). The same data can be replotted to view the inverse correlation between cell size and the amount grown in a specific cell cycle phase or over the entire cell cycle (Figure 3G-I). Similar trends can be seen in the unsmoothed and uninterpolated data (Supplemental Figure S10).

### Cell size predicts timing of G1 exit

Since cell growth during G1 exhibits a near-sizer, we sought to quantify which factors most affect the rate of G1/S progression. We performed multiple-logistic regression using cell size, cell age, and growth rate as predictors of the timing of G1 exit. We found that both cell size (*P* < 10^-10^) and cell age (*P* < 10^-5^) were significant factors, while growth rate was not (*P* > 0.05). Logistic regression with cell volume yields a sharp separation between G1 and S phase cells (Figure 4A), while logistic regression with cell age yields a much shallower slope (Figure 4B). We quantified the accuracy of using cell volume, cell age, or both to predict G1 exit, by calculating the receiver operating characteristic (ROC) curve for all three sets of predictors (Figure 4C). We found that ROC curves obtained from using cell volume or using both cell volume and cell age were nearly identical, with the area under the ROC curve being 0.93 for using cell volume alone and 0.94 when using both cell volume and age. This indicates that while cell age is statistically significant, it is a small factor relative to cell volume in determining the timing G1 exit.

**Figure 4.**
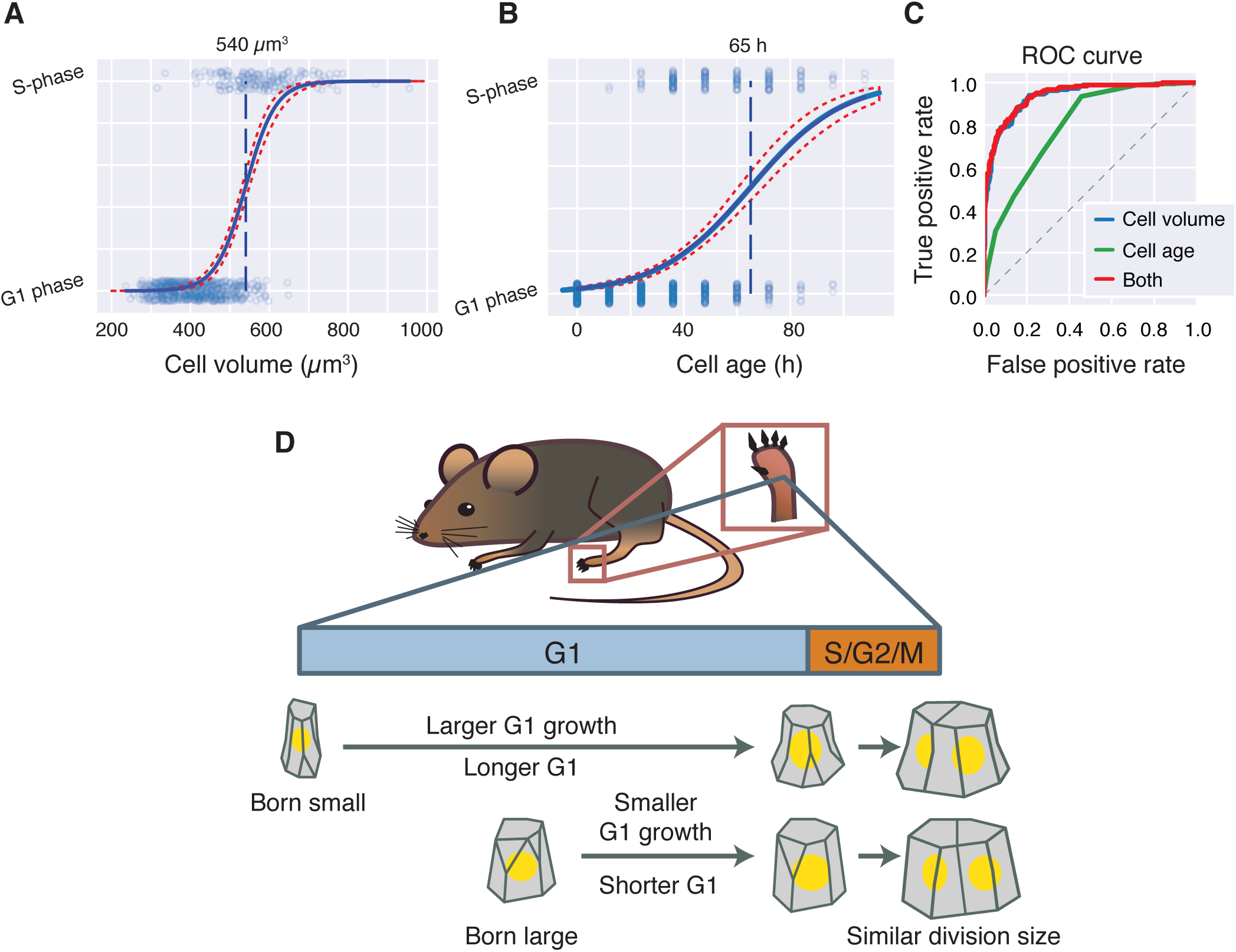
Size-dependent G1/S transition model for *in vivo* cell size control. A-B. The logistic regression predicting the exit from G1 into S phase using cell volume (A) or cell age (B). Dots are individual time-points (*N* = 1,024). Solid blue line is the logistic model, with 95% confidence intervals in dotted red. Dashed blue line is the midpoint. C. The ROC curve from logistic regression models predicting G1 exit using cell volume (blue), cell age (green), or both (red). The dotted line is the null-model of a random predictor. D. Model of skin cell size control *in vivo*. For epidermal stem cells, variation in birth volume are compensated by the size-dependent transition from G1 to S-phase, such that cells born small will spend longer in G1 and grow more in G1. While the control mechanism is insufficiently resolved for S/G2/M, the overall size control is a sizer, where cells divide at a size uncorrelated with their birth size.

## Discussion

Keeping cell size homeostasis during proliferation is crucial for maintaining optimal cell physiology, and may be important for proper tissue geometry and function [26–31]. One way that cell size can be controlled is by coupling cell cycle progression to cell growth through a sizer, in which growth over a single cycle compensates for variation in birth size. This way, all cells reach a similar size at division. However, adders, where cells grow a constant amount over a single cycle independent of birth size, more closely approximate the size control exhibited by a variety of animal cell lines grown *in vitro* [9,10,27,32]. Here, in contrast to expectations from *in vitro* studies, we show that mouse epidermal stem cells exhibit an overall sizer *in vivo*, and variation in cell birth size is compensated for within one cell cycle. Most of this compensation takes place in the G1 phase, but there may also be some compensation during the remainder of the cell cycle (Figure 4D). To our knowledge, this work is the first study examining how cell size control is operating in a mammalian tissue *in vivo*. Furthermore, our work highlights that cell birth size is an important factor in determining the timing of stem cell division.

Our analysis of the growth and division of adult epithelial stem cells *in vivo* has several implications for similar *in vitro* studies. First, the observation of a sizer *in vivo* implies that researchers working on epithelial cells should focus on lines that exhibit a similar sizer phenomenon rather than the wide range of cell lines that do not. Second, there is a dramatic difference between the duration of cell cycle phases *in vitro* and *in vivo*. While the S/G2/M phase of the cell cycle is similar in duration *in vitro* and *in vivo* at ∼12 hours, the G1 phase extends 5-fold from ∼10 hours *in vitro* to ∼50 hours *in vivo*. G1 also accounts for ∼50% of total *in vivo* growth (Supplemental Figure S11), while *in vitro* it is only about ∼25% [10, 12]. Thus, G1 growth and size control is more important *in vivo* than *in vitro*. The G1 sizer we observed may be related to the small size of skin stem cells when they are born (∼350 µm^3^ vs ∼1500 µm^3^ for a HeLa cell) [10]. Fission yeast born with average or smaller size exhibit a sizer, whereas larger-born fission yeast exhibit an adder [2]. Similarly, only small rat basophilic leukemia cells exhibit a sizer [11]. The smallness of stem cells maybe why they exhibit such strong sizer behavior. An alternative explanation for why we observe a sizer *in vivo* but an adder *in vitro* is that cells grow more slowly *in vivo*, with average G1 growth rate being ∼5 µm^3^ hr^-1^ *in vivo* compared to ∼80 µm^3^ hr^-1^ for HeLa cells *in vitro* [10]. A 3-fold change in growth rate clearly impacts size control in *E. coli*, which exhibit an adder when growing quickly but a sizer when growing more slowly [7, 33]. Finally, we note that we have only analyzed skin stem cells and other cell types could have different size control mechanisms *in vivo*.

While our work definitively identifies a G1 sizer *in vivo*, it gives us limited information about the underlying regulatory network. Yeast cells employ the same G1 regulatory network to produce sizers for small-born cells and adders for large-born cells [2,8,34]. This is similarly found in rat leukemia cells [11]. We expect that either of the currently proposed G1 control mechanisms, Rb dilution or p38 activation [35, 36], could theoretically yield sizers or adders depending on parameter values. In addition, an important experimental limitation is the 12h sampling frequency. This low sampling frequency precludes a more careful analysis of growth during S/G2/M phases of the cell cycle. To quantify the effect of poor temporal sampling, we analyzed data produced by [12] of the cell size growth and cell cycle progression of human mammary epithelial cells (HMECs) growing *in vitro* (Supplemental Figure S8A-B). While we saw that size control correlations in G1 were largely unaffected by the low sampling rate, the same was not true for the shorter S/G2/M phases (Supplemental Figure S8C-F). By down-sampling the HMEC dataset to have successively poorer sampling of S/G2/M phases, we observed that a spurious negative correlation between size at G1/S and S/G2/M duration emerges as a result of low temporal sampling, suggesting uncertainty in our estimation of S/G2/M dynamics (Supplemental Figure S8G). Additionally, the low sampling frequency prevents accurate estimation of the growth function, *i.e.*, how rapidly cells grow as a function of their size or cell cycle position. At this point, we can conclude that cell growth is not linear, meaning that cells are not increasing their size at a constant rate through the cell cycle. Rather, larger cells are growing faster than smaller cells, and this relationship between size and growth rate is similar in G1 and S/G2/M cells. However, higher temporal resolution is required to more precisely determine the cell growth function and its relationship to exponential growth (where the cell growth rate is directly proportional to cell size), which has implications for cell size control [9,21,22,37].

We have here analyzed cell growth and division at the level of single cells and neglected potential contributions related to tissue geography. In multicellular tissues, cells experience complex chemical or mechanical feedback from neighboring cells and are regulated by pathways controlling organ size. Certainly, regardless of these multicellular considerations, our observation of a G1 sizer mechanism is inconsistent with a previous model, in which cell growth proceeds linearly while division timing occurs at a constant frequency [20], because cell growth in G1 is clearly coupled to the timing of the G1/S transition. This is also observed in a similar single cell analysis of *in vivo* cell division in the *Arabidopsis* meristem, although the plant stem cells exhibited control mechanism intermediate between sizer and adder [17].

An important open question is how cell autonomous size control is integrated in the context of a multicellular organ. One attractive model is that total cellular growth within a tissue is determined by organ size control mechanisms, and cell size control mechanisms operate at the cell level to quantize that total growth into individual cells. Specifically for the epidermis, local changes in cell density lead to stem cell growth as shown by [13], and this growth is then coordinated with cell division by a size-dependent G1/S transition. This model is consistent with the corpus of studies showing that mutations affecting G1/S control have generally little effect on organ size, but impact cell size and number within the organ [38–40]. Certainly, we anticipate it will be exciting to see if and how specific cell-autonomous size control mechanisms interact with organ size control pathways.

## Methods

### Mice and *in vivo* imaging

All movies used in this study were previously published [13]. Briefly, cells from non-hairy plantar skin were imaged with two-photon microscopy every 12h for 4 to 7 days. Regions were re-visited in independent imaging sessions using micro-tattoo fiducial marks for orientation and intrinsic idiosyncratic features of each skin region for alignment. In between imaging time-points, mice were returned to normal activity. Three datasets used in this study were of mice expressing *CMV-mKO2-hCdt1(30/120)* [41], *K14-actin-GFP* [42], and *K14-H2B-Cerulean* [13], where *K14* is the human *K14* promoter expressed only within the keratinocyte lineage. Two regions were used from Mouse 1, which was imaged for 7 days (168 h). A third region from Mouse 2 was imaged for 4 days (96 h). Data acquired from both animals were similar (Supplemental Figure S2), although the shorter movie duration in Mouse 2 limited our sampling of longer cell cycles in Region 3.

### Cell volume reconstruction and cell cycle annotation

All cell volumes were manually segmented in FIJI using PolygonRois [43]. Automatically tracked lineages from Mesa, Kawaguchi, & Cockburn, *et al*. (2018) were filtered for cells that are born and divide within the duration of the movie [13]. A custom FIJI applet was built to display automatically tracked cell centroids on the movie to facilitate manual segmentation. Mistakes made by the automatic tracking were fixed manually. Cell volume was calculated by summing the cell area throughout all z-slices and multiplying by the z-step size. Custom Jpython scripts were used to automate ROI measurement and export in FIJI. Repeated volume reconstructions were made to quantify intra-user variation in segmentation. Repeat growth curves were generated for 8 cells (*N* = 48 time-points) chosen at random and the agreement between the original and the repeat measurements were quantified (Supplemental Figure 3I-J). Daughter cell volumes were quantified in the same way as mother cell volumes. Cells whose daughter volumes could not be estimated were excluded from the analysis involving interpolated division volumes (*N* = 30). For comparison of cell volume to cross-sectional cell area, the z-position of the cross-section was determined by [13] to be the z-position corresponding to the largest projected nuclear area.

Cell cycle transitions were annotated manually based on the fluorescent FUCCI-G1 reporter mKO2-hCdt1(30/120). Because the illumination was not constant throughout the movie, the total intensity of the G1 reporter was not comparable across time-points. Therefore, the G1 exit frame was annotated as the frame at which the G1 reporter within the cell nucleus became indistinguishable from the local background (Supplemental Figure S1). The FUCCI-G1 reporter also had variable expression in the basal layer cells so that a subset of cells never had visible expression throughout their cell cycle. These cells were excluded from the analysis. Mitotic cells were manually identified by their rounded cell shapes and visible nuclear envelope breakdown (Supplemental Figure S7H). The fidelity of the segmentation and tracking was assessed by examining the 3D segmentation overlaid on the original movie images (**Supplemental Movie S1-2**).

### Volume growth analysis

Cell volume growth curves and cell cycle annotations were collated in Python using pandas [44]. We fitted cell volume growth curves to smoothing cubic splines using numpy and scipy [45, 46]. A high smoothing factor (10^5^) was used to ensure relatively stiff spline fits (Supplemental Figure S3A-F). We used the smoothed data for analysis because we expect volume growth to be both smooth and generally to be monotonically increasing through the cell cycle. Error in individual time-points was estimated as the magnitude of residuals (absolute difference between volume time-points and fitted curves) (Supplemental Figure S3H). Growth curves with fewer than 4 points were kept as unsmoothed. Volume growth curves were also fitted to linear and exponential models and fits were compared using their residuals. Growth rates were estimated from the smoothed growth curves using backwards difference (window size = 1). Confidence intervals for CVs and P-values for differences in CVs were calculated from bootstrap analyses.

### Nuclear volume reconstruction

To quantify nuclear volume, the H2B-Cerulean channel was automatically segmented by thresholding pixels that were above the 50th percentile in intensity within a 31-pixel local neighborhood using the scikit-image module in Python (Supplemental Figure S5) [47]. Small holes and small objects in the thresholded nuclear masks removed by binary erosion and dilation. The nuclear volume was calculated as the number of above-threshold pixels within the manual cell outline segmentation, multiplied by the pixel:µm conversion ratio and z-step size. Nuclear volumes calculated for cells in mitosis were discarded because of nuclear envelope breakdown.

### Analysis of framerate limitations

We analyzed *in vitro* human mammary epithelial cell (HMEC) growth data from [12], who acquired time-lapse phase and fluorescence images of asynchronously growing and dividing cells expressing fluorescent reporters of cell size and cell cycle phase marking the G1/S transition. We examined the time-series from these data, which were taken at 10 minute time intervals. HMECs in these conditions had a ∼20 hour cell division cycle and exhibited substantial size control at the G1/S transition, but not during S/G2/M phases. The data were down-sampled to quantify how lowering temporal resolution can affect cell size control correlations. The data decimation factor was determined by down-grading the time resolution in [12] until it had the same average number of frames per cell cycle as we have analyzed here for epidermal stem cells *in vivo* (Figure 2C; Supplemental Figure S8A-B). For each cell, we down-sampled its time-series starting with a randomly selected time-point and the process was repeated 500 times. The resulting randomized distributions of correlations were compared to the correlations in the original data (Supplemental Figure S8C-E). While the G1 size control correlations are similar between original and down-sampled data, growth during S/G2 is systematically underestimated and a larger error in the correlation between G1 exit size and growth during S/G2 is introduced (Supplemental Figure S8F). To further examine the effect of low temporal resolution on S/G2 dynamics, we used successively larger data decimation factors and observed that a spurious negative correlation between size at G1/S and S/G2 duration could result from poor temporal sampling (Supplemental Figure S8G).

### Statistical analysis

All statistical analyses were done in Python using numpy/scipy and statsmodels [48]. Multivariate regression analysis was done using OLS and Logit from statsmodels. All correlation coefficients (*R*) reported in the text are Pearson’s correlation. *P*-values reported for Pearson’s correlation are tested against the null-hypothesis of *R* = 0. All intervals are given in the text as mean ± std, unless otherwise noted. All code used in the study is available at https://github.com/xies/mouse_skin_size_control/.

## Supporting information

Supplemental Movie S1. Tracking epidermal stem cells through 7 days of growth and division.

Supplemental Movie S2. 3D reconstruction of epidermal stem cells.

## Acknowledgements

We thank members of the Skotheim lab, Evgeny Zatulovskiy, Daniel Berensen, Kurt Schmoller, Clotilde Cadart, for discussion and comments on the manuscript. We thank Valentina Greco and Katie Cockburn for discussions, comments, and facilitating our understanding of the data from Mesa et al. (2018). This work was supported by the NIH through R01 GM115479 and F32 GM129878 (XS).

---

**Supplemental Figure S1.**
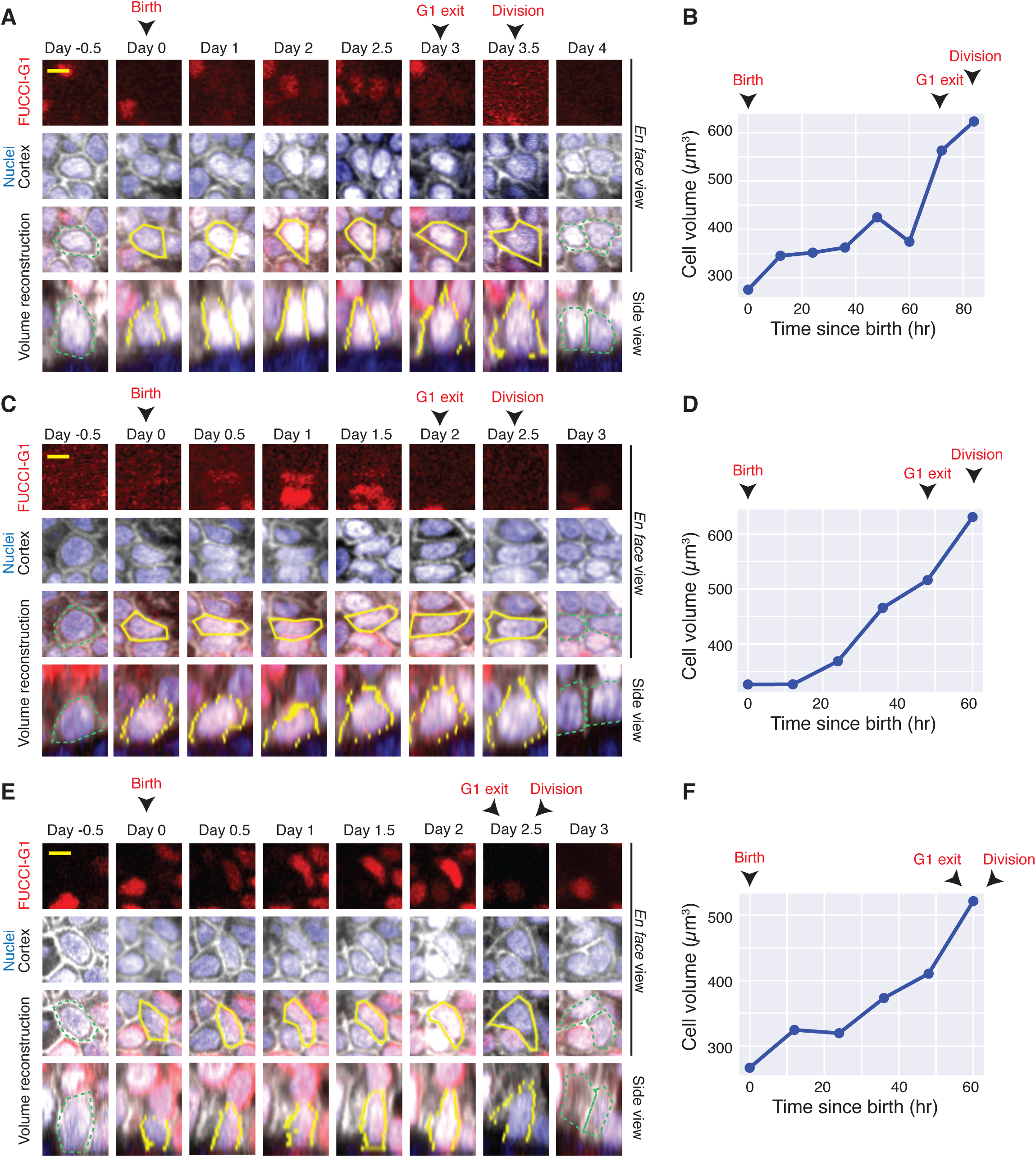
Examples of 3D reconstruction of epidermal stem cells. A-F. Three examples of the volume reconstruction of single epidermal stem cells. The FUCCI-G1 reporter is shown in red, the nucleus in blue, and the actin cortex in gray. The manually segmented cell outlines are shown in yellow on top of the merged images showing *en face* and side views. The cell cycle landmarks birth, G1 exit, and division are annotated. The parent cell and daughter cells are outlined in dotted green. The volume growth curve is quantified for the cells shown in their respective neighboring panels. Scale bars are 5 µm. Related to Figure 2.

**Supplemental Figure S2.**
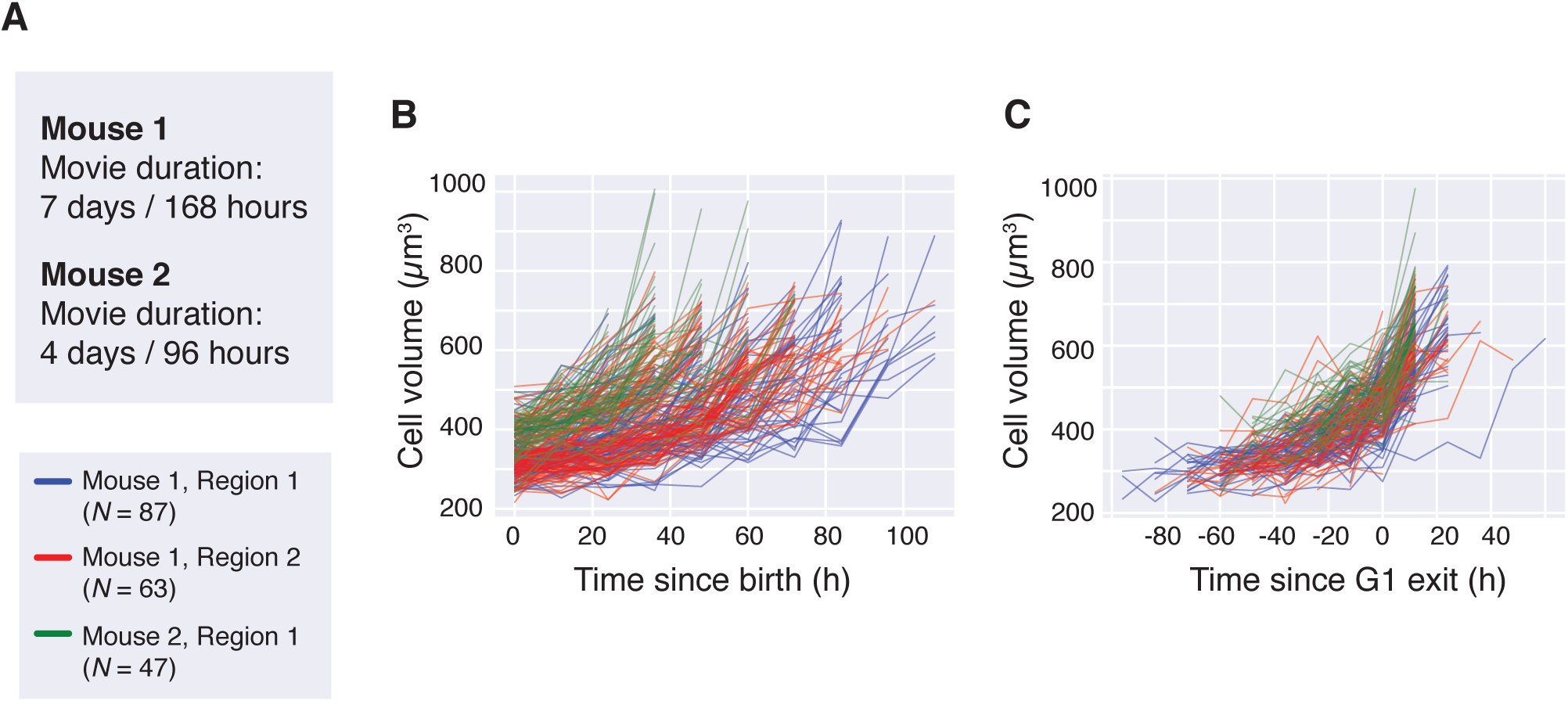
Epidermal stem cell growth dynamics *in vivo*. A. Summary of the multiple skin regions used in this study. Two regions were analyzed from mouse 1, which was imaged for 7 days. One region was analyzed from mouse 2, which was imaged for 4 days. B. Volume growth curves aligned by birth (*N* = 197). Each independent region is as indicated in (A). C. Volume growth curves aligned by G1 exit. Related to Figure 2.

**Supplemental Figure S3.**
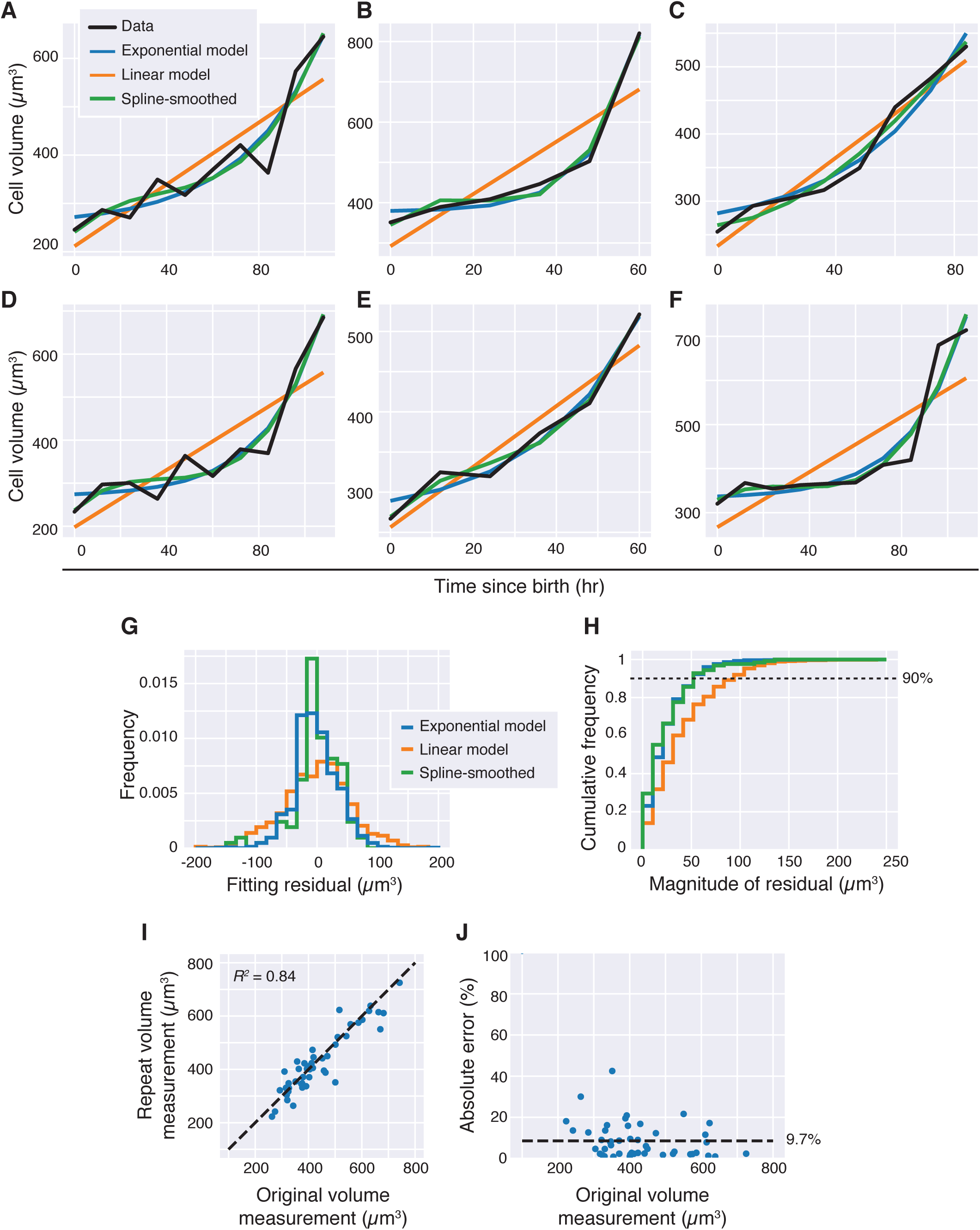
Cell growth curves and segmentation noise. A-F. Examples of individual volume growth curves. The fits to linear (orange), exponential (blue), and cubic smoothing spline (green) models are shown. The empirical data is shown in black. G. The distribution of fitting residuals from linear, exponential, and smoothing spline fits (*N* = 1,159). H. The cumulative distribution of the magnitude of residuals. The dotted line denotes the 90% cumulative probability. I. Intra-user variation in cell volume reconstruction for 48 independent time-points. The dotted line is the identity. J. The intra-user absolute error rate, defined as the absolute difference between repeat and original measurements normalized by the original volume measurement. The average error of 9.7% is shown as a dotted line. Related to Figure 2.

**Supplemental Figure S4.**
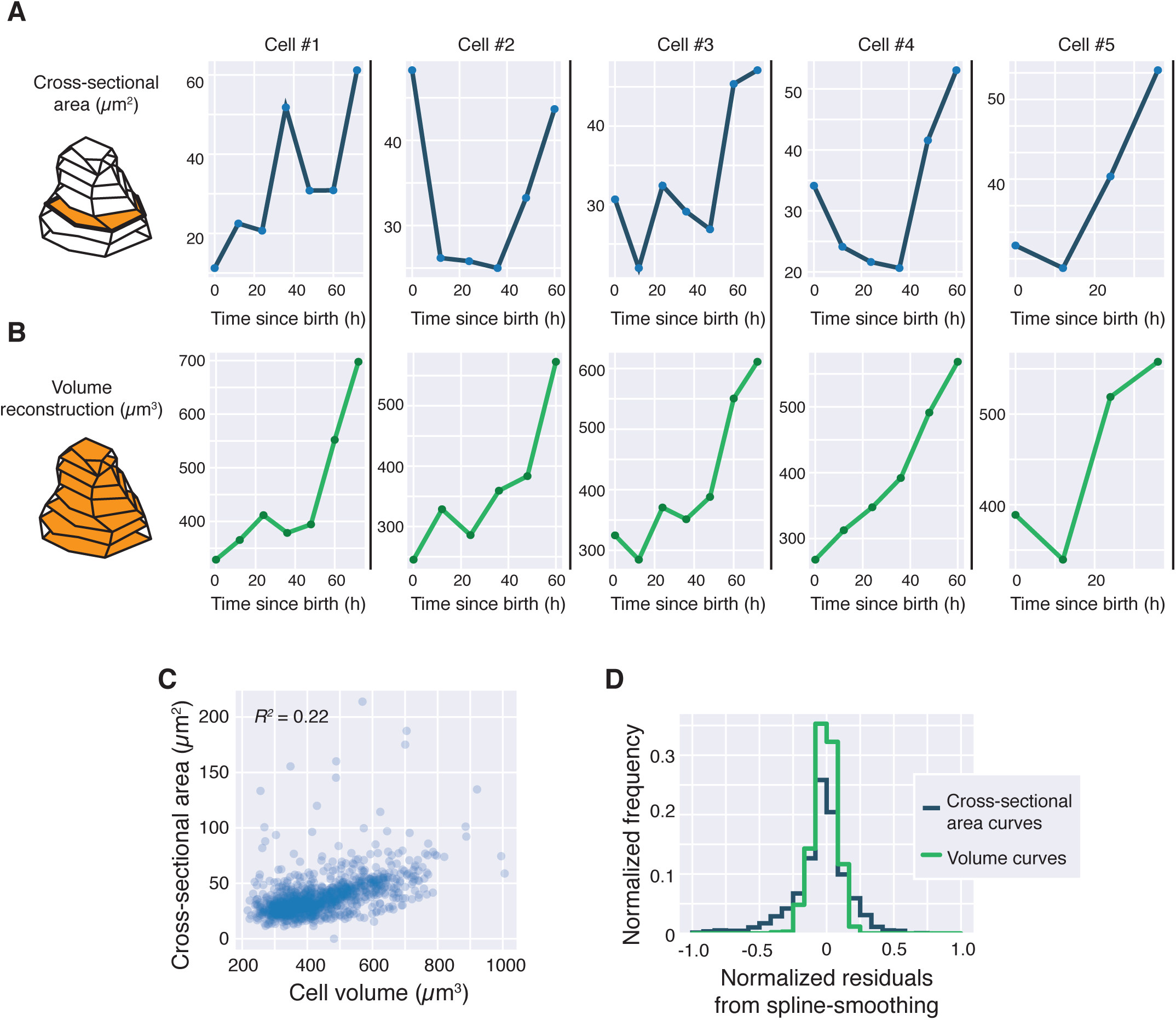
Comparison of cross-sectional area and cell volume as estimates for cell size. A. Five representative growth curves of the cross-sectional cell area through an entire cell cycle. B. The cell volume growth curves for the same five cells shown in (A). C. The correlation between cross-sectional area and cell volume (*N* = 827). D. The magnitudes of the normalized residuals (residuals normalized by the empirical data) from fitting area growth curves (blue) or volume growth curves (green) to smoothing splines. The magnitudes of normalized residuals are larger for cross-sectional area measurements than for volume measurements (*P* < 10^-15^, one-tailed two-sample KS-test). Related to Figure 2.

**Supplemental Figure S5.**
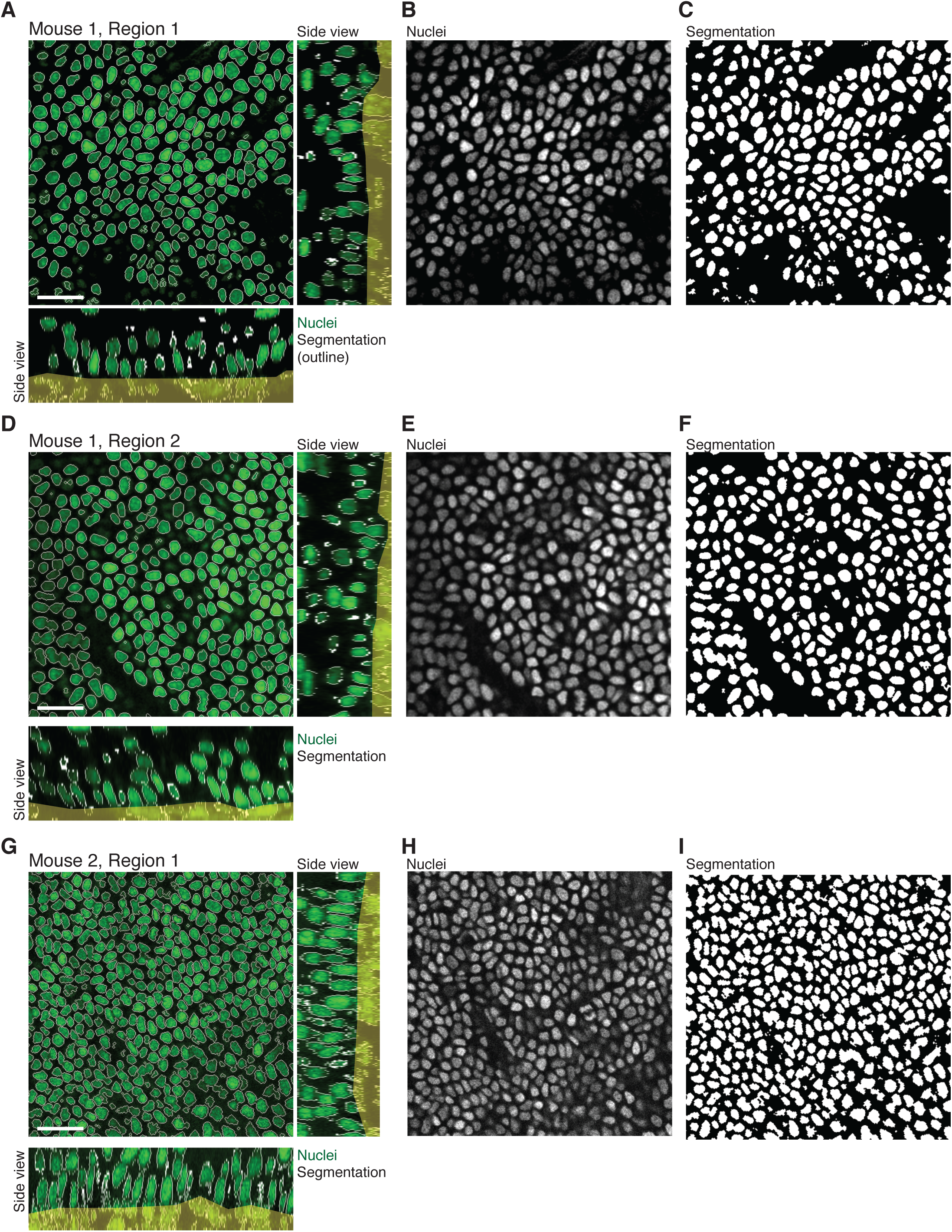
Nuclear segmentation for nuclear volume reconstruction. A, D, G. The results of automatic nuclear segmentation for the three independent regions. The H2B-Cerulean nuclear reporter is shown in green. The thresholded outlines are shown in white. *En face* views and side views are shown. The dermis underneath the basal layer (non-epidermal tissue) is masked out in yellow. B, E, H. The H2B-Cerulean nuclear reporter images. C, F, I. The nuclear masks generated by automatic thresholding of the H2B-Cerulean signal. Related to Figure 2.

**Supplemental Figure S6.**
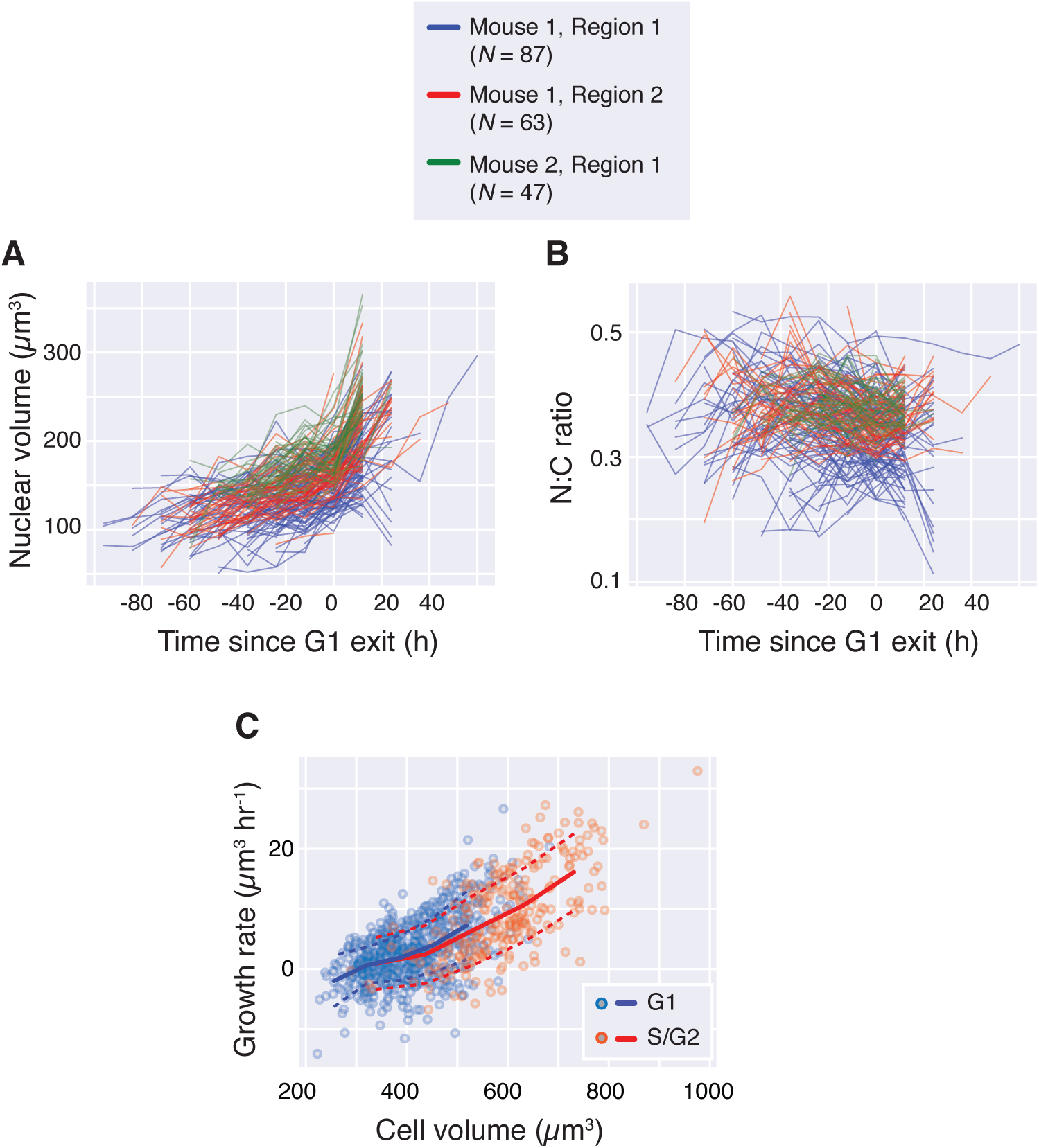
Epidermal stem cell nuclear volume, N:C ratio, and volume growth rate. A. The nuclear volumes aligned by G1 exit (*N* = 197). Each imaging region is as indicated. B. The N:C volume ratios are plotted with respect to time and aligned by G1 exit. C. The volume growth rate (backwards differences) is plotted against cell volume, both estimated from unsmoothed data (*N* = 946). Blue dots denote G1 cells, orange dotes denote S/G2 cells. Binned means are shown in blue (G1) and orange (S/G2). Dotted lines show standard deviations. Related to Figure 2.

**Supplemental Figure S7.**
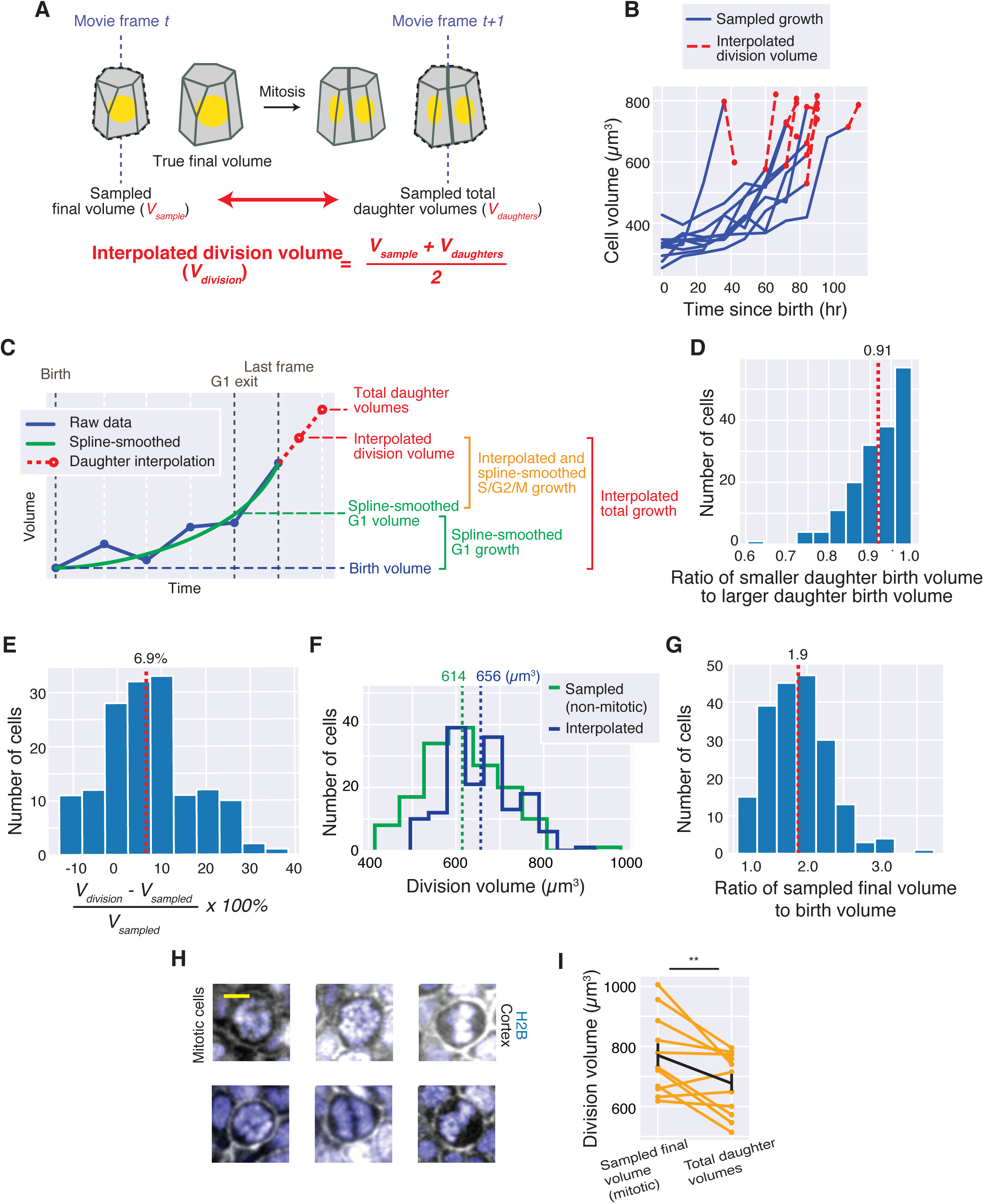
Estimating final cell division volume. A. Schematic of interpolated division volume estimation. We took the average between the sampled final volume and the sum of the volume of the two daughter cells. B. The cell volume growth curves are shown for ten representative cells, where the interpolated division volume is shown in red. C. Schematic of cell volume estimations used in Figure 3. We fit the data to a smoothing spline to estimate G1 exit volume. We use the interpolated division volume as defined in (A). D. The ratio of the birth volume of the smaller daughter cell to its larger sister cell (*N* = 167). Dotted line denotes the median. E. The distribution of the difference between sampled and interpolated division volumes, normalized by the sampled volume. Dotted line denotes the mean. F. The distribution of the sampled final volume (green) and the interpolated division volume (blue) are shown for cells whose last frame is not during mitosis. Dotted lines denote the means for the distribution of corresponding color. G. The distribution of the ratio between the sampled final volume to birth volume. Dotted line denotes the mean. H. Examples of mitotic cells. The histone reporter is shown in blue. The actin cortex is shown in gray. Scalebar is 5µm. I. The differences between the sampled final volume and the interpolated division volume are shown for cells that are in mitosis in their last frame (*N* = 11). Black line denotes the mean. Cells in mitosis are larger than the sum of the two daughter cell volumes. (**: one-tailed paired T-test, *P* < 0.01). Related to **Figure 2, 3**.

**Supplemental Figure S8.**
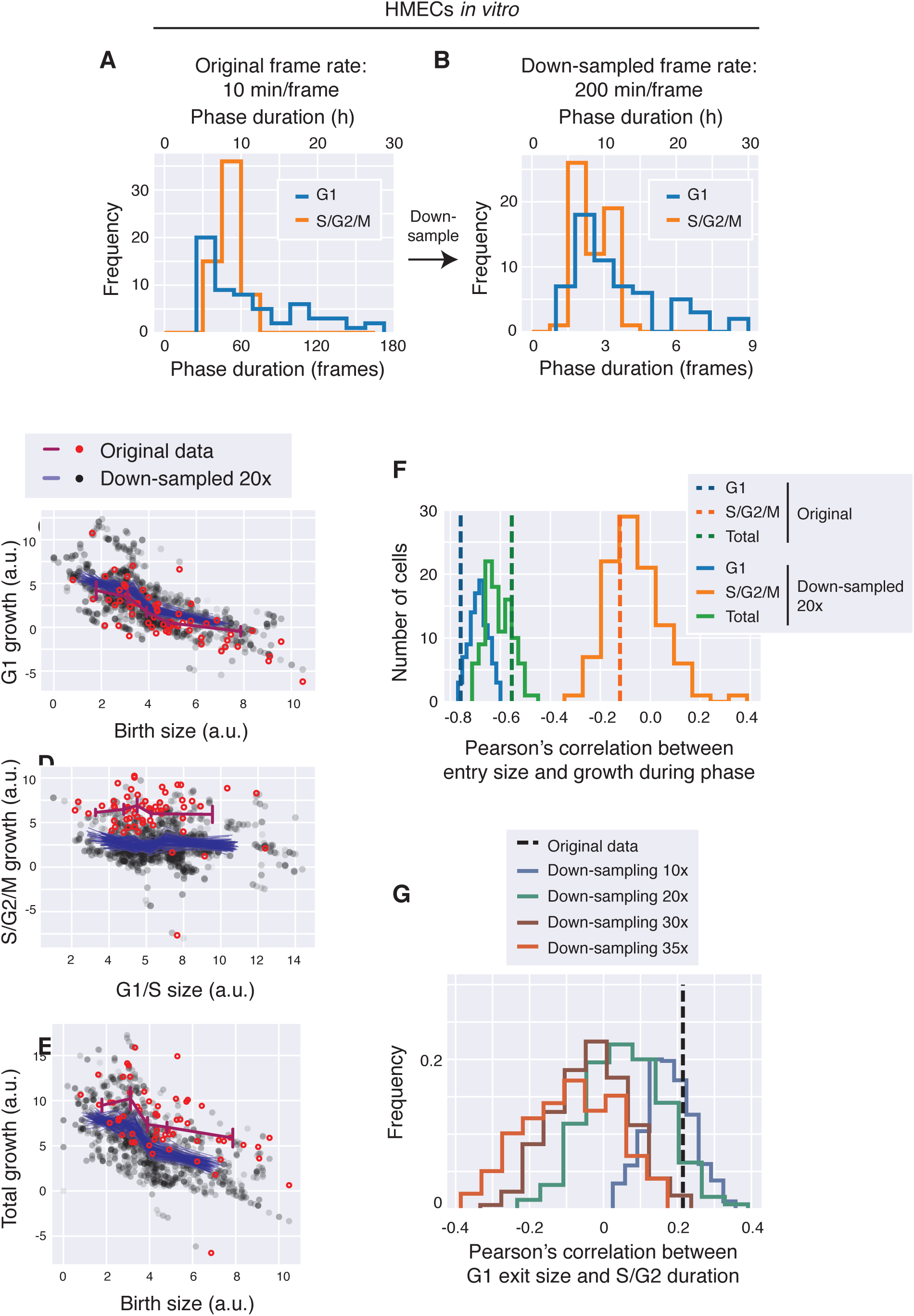
Estimating the effect of low temporal resolution on size control correlations. A. The distribution of cell cycle phase durations of HMECs growing *in vitro* from [12]. The corresponding number of frames is shown. B. The data in (A) were down-sampled 20-fold so that we had the same number of time points as the *in vivo* data. Compare with Figure 2C. C-E. Cell size control correlations are shown for HMECs comparing original and down-sampled data. Original data are shown in red, while the ensemble of 500 randomly down-sampled data are shown in black. The solid lines denote binned mean data for original (red) or down-sampled (blue) data. Cell size is measured from the cell size reporter generated by [12]. Birth size is plotted against G1 growth in (C). Cell size at G1/S is plotted against S/G2 growth in (D). Birth size is plotted against total growth in (E). F. The Pearson’s correlation (*R*) between the cell size at the indicated phase-entry and growth during the indicated phase are shown for G1 (blue), S/G2 (orange), and the total cell cycle (green). The distribution of *R* from randomly down-sampled data are shown in solid lines. Dotted lines denote the value of *R* in the original data for their corresponding colors. *R* for S/G2 phase in particular shows a broader distribution in the down-sampled data. G. The Pearson’s correlation (*R*) between G1 exit size and S/G2 duration for increasing down-sampling factors. Dotted line indicates the *R* in the original data. As the temporal sampling becomes poorer, the distribution of *R* shifts left, indicating a spurious negative correlation may be introduced by poor temporal sampling. Related to Figure 3.

**Supplemental Figure S9.**
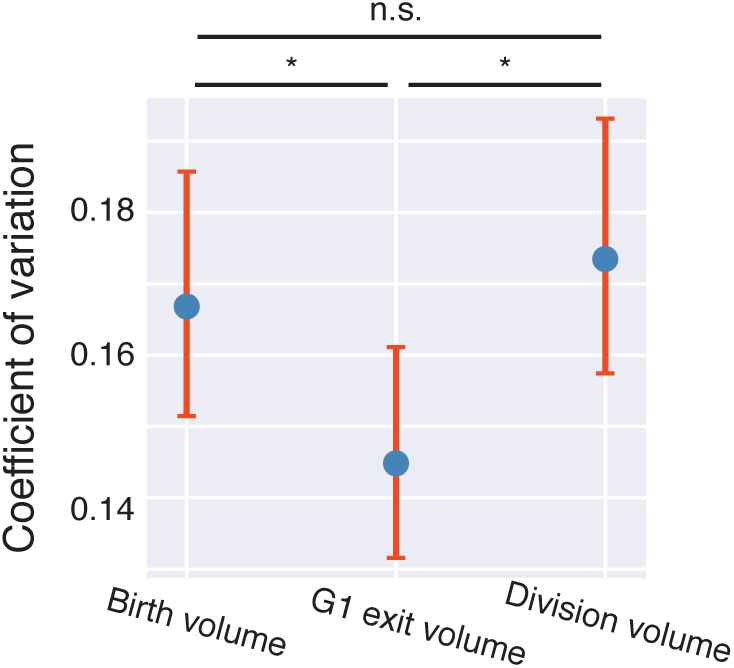
The CV of cell volume at cell birth, G1 exit, and cell division. *N* = 197. Error bars show the 95% confidence interval estimated using bootstrapping. (*: *P* < 0.05, n.s.: *P* > 0.5; one-tailed bootstrap test).

**Supplemental Figure S10.**
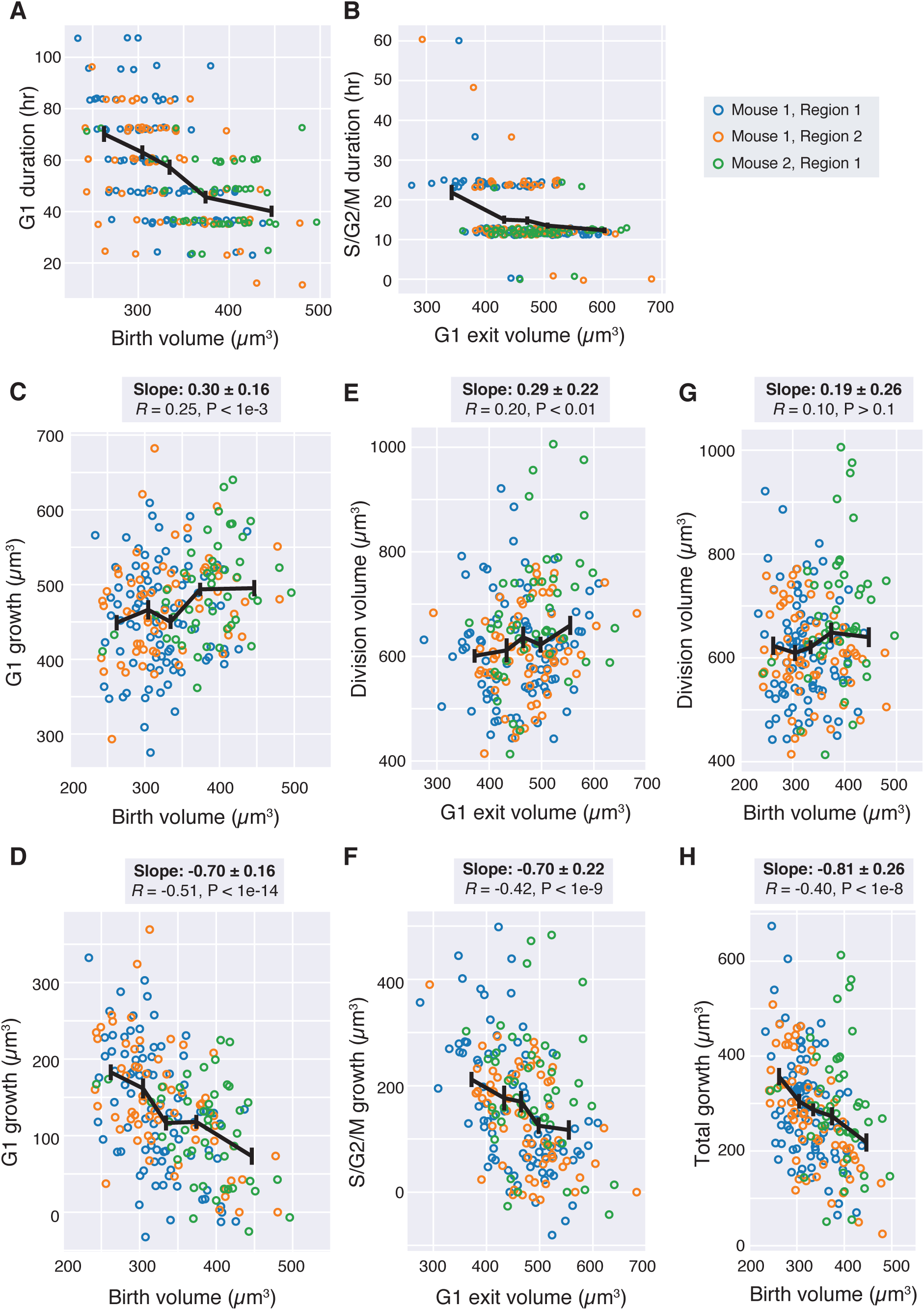
Cell size control correlations showing unsmoothed data from individual regions. A. Cell birth volume is plotted against the duration of the G1 phase. Cells from each individual region are shown in different colors. B. The volume at which cells exit G1 is plotted against the duration of S/G2/M phases. C-D. Birth volume is plotted against the volume at G1 exit (C) or the amount of growth during G1 (D). E-F. The G1 exit volume is plotted against the volume at division (E) or the amount of growth during S/G2/M (F). G-H. The birth volume is plotted against the division volume (G) or the total amount of growth (H). All data are unsmoothed and contain no daughter volume interpolation. Black lines show binned means ± SEM. Slopes reported correspond to the slope of the linear regression and 95% confidence intervals. *R* values are Pearson’s correlations, with *P*-values reported against the null-hypothesis of *R* = 0. We note that not using smoothed data leads to the same qualitative conclusions as using smoothed data. Related to Figure 3.

**Supplemental Figure S11.**
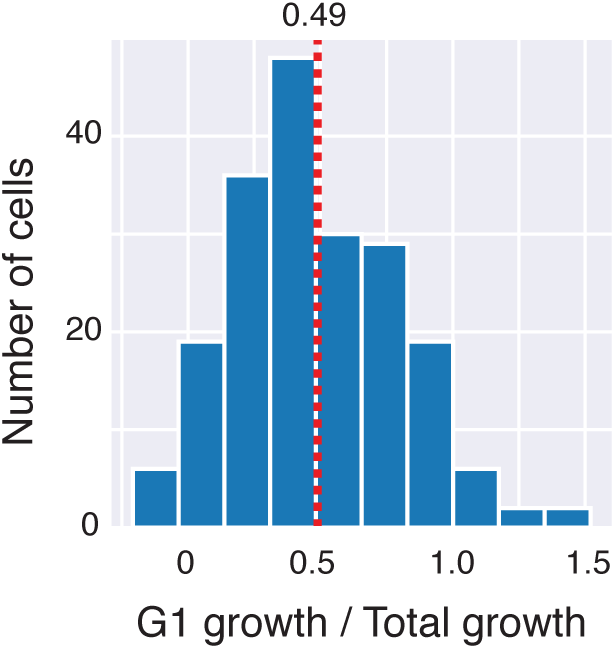
The distribution of the ratio between growth during G1 over total growth. *N* = 167, dotted line denotes the mean.

**Supplemental Movie S1. Tracking epidermal stem cells through 7 days of growth and division.**

A single z-slice through time from the time-lapsed *in vivo* 4D imaging of mouse epidermal stem cells. Actin-GFP (cell cortex) is shown in gray, H2B-Cerulean (nucleus) is shown in blue.

Manually segmented cell outlines are shown in yellow. The cell’s identifying index is overlaid on the cell, demonstrating the fidelity of cell tracking.

**Supplemental Movie S2. 3D reconstruction of epidermal stem cells.**

A single time-frame from the time-lapsed *in vivo* 4D imaging of mouse epidermal stem cells. Z-slices are shown from apical (where differentiated cells reside) to basal slices (where stem cells reside). Actin-GFP (cell cortex) is shown in gray, H2B-Cerulean (nucleus) is shown in blue. Manually segmented cell outlines are shown in yellow. The cell’s identifying index is overlaid on the cell.

